# Transcriptomic analysis of insecticide resistance in the lymphatic filariasis vector *Culex quinquefasciatus*

**DOI:** 10.1101/589028

**Authors:** Walter Fabricio Silva Martins, Craig Stephen Wilding, Alison Taylor Isaacs, Emily Joy Rippon, Karine Megy, Martin James Donnelly

**Author notes:** These authors contributed equally to this work (Walter Fabricio Silva Martins, Craig Stephen Wilding).

## Abstract

*Culex quinquefasciatus* plays an important role in transmission of vector-borne diseases of public health importance, including lymphatic filariasis (LF), as well as many arboviral diseases. Currently, efforts to tackle *C. quinquefasciatus* vectored diseases are based on either mass drug administration (MDA) for LF, or insecticide-based interventions. Widespread and intensive insecticide usage has resulted in increased resistance in mosquito vectors, including *C. quinquefasciatus*. Herein, the transcriptome profile of Ugandan bendiocarb-resistant *C. quinquefasciatus* was explored to identify candidate genes associated with insecticide resistance. Resistance to bendiocarb in exposed mosquitoes was marked, with 2.04% mortality following 1h exposure and 58.02% after 4h. Genotyping of the G119S *Ace-1* target site mutation detected a highly significant association (*p*<0.0001; OR=25) between resistance and *Ace1*-119S. However, synergist assays using the P450 inhibitor PBO or the esterase inhibitor TPP resulted in markedly increased mortality (to ≈80%), suggesting a role of metabolic resistance in the resistance phenotype. Using a novel, custom 60K whole-transcriptome microarray 16 genes significantly overexpressed in resistant mosquitoes were detected, with the P450 *Cyp6z18* showing the highest differential gene expression (>8-fold increase vs unexposed controls). These results provide evidence that bendiocarb-resistance in Ugandan *C. quinquefasciatus* is mediated by both target-site mechanisms and over-expression of detoxification enzymes.

## INTRODUCTION

Lymphatic filariasis (LF), is a major cause of chronic and permanent disability in tropical and subtropical regions as a result of lymphoedema, elephantiasis and hydrocele ^1,2^ and is endemic in 83 countries with more than 1.2 billion people at risk of infection, especially in Southeast Asia and Africa ^3,4^. In sub-Saharan Africa the causal agent of LF is the nematode *Wucheraria bancrofti*, which can be transmitted by both Culicine and Anopheline mosquitoes with *Culex quinquefasciatus* the major vector in urban settings in East Africa ^5^.

In contrast to other vector-borne disease control programmes such as malaria and dengue that use anti-vector interventions as the major strategy, the Global Program to Eliminate LF (GPELF) is based on Mass Drug Administration (MDA) of anthelmintics to reduce *W. bancrofti* transmission. Nevertheless, vector control is recommended as an intervention for LF eradication in regions where successful implementation of MDA is challenging, for instance in very remote areas, or where LF is co-endemic with loiasis which can result in adverse reactions to the drug cocktail used for MDA ^4,6^. Modelling and field studies have shown that integration of vector control into MDA programmes can reduce the required number of chemotherapy rounds and consequently the time frame to achieve the microfilaria (MF) prevalence threshold necessary for successful interruption of LF transmission ^1,7^.

Although vector control has successfully reduced the burden of vector-borne diseases worldwide, the recurrent and extensive application of insecticides in endemic regions has also triggered an increase in the level of insensitivity to those insecticides approved for public health ^8,9^. In addition, the limited number of insecticides available and the occurrence of cross-resistance between different classes is especially worrying for the sustainability of vector control ^10,11^. Consequently, identification and monitoring of resistance patterns, and understanding the underlying mechanisms is crucial for extending the lifespan of currently available insecticides, as well as for planning more effective vector control programmes.

Insensitivity to insecticides in arthropods is thought to result mainly through mutations in target-site genes and/or overproduction of detoxifying enzymes ^12,13^. Susceptibility studies in *C. quinquefasciatus* from diverse geographical regions have associated two main target-site mutations to resistant phenotypes. The L1014F mutation in the voltage-gated sodium channel gene, conferring *kdr* (knockdown resistance), has been associated with pyrethroid and DDT resistance, whilst the G119S mutation in the acetylcholinesterase (*Ace-1*) gene is linked to resistance to carbamates and organophosphates ^13–16^. Metabolic resistance, which involves the over-expression, or increased catalytic capability of metabolic enzymes, is a less tractable mechanism since members of diverse gene families including carboxy/cholinesterases, glutathione S-transferases (GSTs) and cytochrome P450 monooxygenases (P450s) have previously been associated with resistance to different classes of insecticide in a range of vector species ^17–19^. Over-expression of detoxification genes can be triggered by a range of mechanisms including gene duplication ^20^, as observed for the resistance to organophosphates in *C. quinquefasciatus* mediated by esterases ^21^, *cis*-regulatory elements ^22,23^, *trans*-regulatory elements, or changes in post-transcriptional repression due to differential expression of miRNAs ^24^.

Due to the diversity of genes or gene families involved in metabolic resistance, identification of candidate genes requires an agnostic survey of the patterns of gene expression associated with resistant phenotypes. Recently, studies have applied either microarray or RNA-seq platforms to elucidate the relationship between gene expression and insecticide resistance ^25–27^ although to date, most of these whole-transcriptome studies in vector insects are restricted to mosquitoes of the genus *Anopheles*. Despite the role of *Culex* in transmission of several pathogens such as filarial worms and West Nile virus (WNV) ^28^, and reports of high levels of insecticide resistance, few studies have addressed the relative impact of metabolic resistance in *C. quinquefasciatus* ^29,30^ particularly at a whole-transcriptome scale (although see ^31,32^).

In addition to their role as disease vectors, *C. quinquefasciatus* are nuisance biters and failure to effectively control this species can lead to the perception of malaria control failure which may ultimately lead to the rejection of controls (e.g. IRS and ITNs) ^15,33^. In this study, we report the results of *C. quinquefasciatus* susceptibility bioassays for six insecticides (DDT, permethrin, deltamethrin, bendiocarb, fenitrothion and lambda-cyhalothrin) from Nagongera, Tororo District, Uganda. We then report and apply a novel 8×60K whole-transcriptome microarray to identify candidate genes associated with bendiocarb insecticide resistance, the active ingredient used in IRS control in Uganda at the time of collection ^34^.

## MATERIALS AND METHODS

### Sample collection

Mosquitoes were collected in Nagongera, Tororo, Uganda (0° 47’ 48.9978", 33° 58’ 47.1") between June and July 2012. Resting adult *C. quinquefasciatus* were collected exclusively inside houses using aspirators and transported to the insectary. From these collections, blood-fed females were maintained in individual Eppendorf tubes lined with moist filter paper to encourage egg laying ^35^. From these, 64 females laid at least one egg batch. Egg batches were floated in water-filled trays and emergent larvae fed on Tetramin fish food. Pupae were transferred to cages and adults allowed to emerge. Eclosion cages were changed every 3 days so that cages contained only 3-5 day-old adults. Adult mosquitoes were fed *ad libitum* on 10% glucose solution and used for insecticide susceptibility testing. Genomic DNA from each female from which egg rafts were obtained to found the colony was individually isolated using a DNeasy kit (Qiagen) then used for identification of *C. quinquefasciatus* using a diagnostic PCR assay ^36^. In addition to these field-collected mosquitoes, a laboratory colony of *C. quinquefasciatus* from the Tropical Pesticides Research Institute (TPRI) Tanzania was used as a susceptible reference strain ^37^ for the microarray study.

### Insecticide susceptibility test

Bioassays were performed using test kits and insecticide-impregnated papers according to standard WHO methods (WHO) ^38^ with adult F1 3-5 day-old females assayed in four replicates of 25 non-blood fed mosquitoes. Tests were performed with papers impregnated with diagnostic concentration for six insecticides: DDT (4%), permethrin (0.75%), fenitrothion (1%), lambda-cyhalothrin (0.05%), bendiocarb (0.1%) and deltamethrin (0.05%). Mosquitoes were exposed for 1h, with the exception of bendiocarb and deltamethrin where, following the results of initial 1h exposures, additional four-hour exposures were used to increase the discrimination between resistant and unexposed sympatric mosquitoes. Control assays were performed with 25 mosquitoes exposed to non-insecticide treated papers.

Insecticide exposed mosquitoes were then transferred to clean holding tubes and provided with 10% glucose for a 24-hour period after which mortality was recorded with dead (susceptible) mosquitoes collected and individually stored on silica gel whilst alive (resistant) mosquitoes had a hind leg removed (stored on silica) and the whole body stored in RNAlater (Sigma Aldrich). RNAlater stored mosquitoes were initially held overnight at 4° C to allow the solution to penetrate the carcass before storage at −20° C until RNA isolation.

### Synergist assays

Synergist tests were carried out following the procedure described above with an additional pre-exposure to three synergist compounds to investigate the potential mechanisms of metabolic resistance for bendiocarb and deltamethrin insecticides. For each synergist assay, four batches of 20-25 mosquitoes were either pre-exposed to impregnated papers (12cm×15 cm, Whatman grade no.1 filter paper) with 4% PBO (piperonyl butoxide – CYP450 synergist), 10% TPP (triphenyl phosphate – esterase synergist) or 8% DEM (diethyl maleate – GST synergist) for one hour, then exposed to bendiocarb (0.1%) or deltamethrin (0.05%) for four hours, followed by a 24h recovery period. Synergist only controls were run simultaneously.

### *Ace-1* genotyping of bendiocarb phenotyped mosquitoes

Prior to microarray analysis all mosquitoes were genotyped for the G119S mutation in *Ace-1*. DNA was isolated from the amputated hind leg of resistant mosquitoes with 50 μl of 10% Chelex 100 and 2 μl of proteinase K (10 mg/mL). The homogenate was incubated at 94°C for 30 min followed by centrifugation at 6,000 rpm for 10 min to collect the supernatant.

A *Taq*Man assay, designed to detect the G119S mutation in the acetylcholinesterase gene ^39^, was utilised for genotyping with reaction mixtures composed of 1 μl of genomic DNA, 1x SensiMix II probe, 400 nM of each primer and 100 nM of each probe in a final volume of 50 μl. Thermocycling was performed on the Stratagene MX3005P and consisted of 95°C for 10 min and 40 cycles of 92°C for 15sec and 60°C for 1 min with endpoint discrimination.

### Microarray

#### 8 X 60K array construction and study design

The pattern of gene expression of bendiocarb resistant mosquitoes was investigated using a custom designed *C. quinquefasciatus* whole genome oligonucleotide microarray (Agilent ID 039759). An 8 × 60K microarray format with 60-mer oligonucleotide probes was designed to cover a variety of targets using eArray (http://earray.chem.agilent.com/earray/). The majority of this array encompassed three probe replicates (56,598 probes) for each of the 19,018 transcripts in the CpipJ1.3 gene build ^40^. Additionally, we downloaded 205,396 ESTs from VectorBase. From these, we identified 1,987 contigs and 4,109 singleton sequences and designed a single probe for each (from these 1,987 contigs, only 1,935 had probes successfully designed and from 4,109 singletons 2,862 had successful probes designed). Additionally, three probe replicates were designed to each of four alternative GST transcripts not annotated in VectorBase ^41^, and 25 additional replicated probe groups (10 replicates) to allow estimation of reproducibility (CV (coefficient of variation) probes) (Fig. 1A). Full details of the array design are given in ArrayExpress (http://www.ebi.ac.uk/arrayexpress/) with the accession number A-MTAB-649.

From the six insecticide susceptibility test groups, we chose bendiocarb selected mosquitoes for the microarray analysis since we observed only moderate mortality (58.02%, 95% CI 51.53-64.25%) for this insecticide following 4h exposure (Fig. 2) thereby allowing clear discrimination between resistant and unexposed sympatric mosquitoes and we also detected an increase in mortality in the presence of two synergists (See Fig. 2; TPP and PBO), indicating the likely involvement of metabolic resistance. Three experimental conditions were employed: Tororo_Resistant (following 4h bendiocarb exposure), Tororo_Control (4h exposure to control papers) and TPRI_Control (4h exposure to control papers). All mosquitoes were wildtype 119G homozygotes for *Ace_1*. Comparison between the bendiocarb selected samples and controls (Tororo_Control and TPRI_Control) was performed on three RNA pools per group using an interwoven loop design (Fig. 1B) as described by Vinciotti *et al*. ^42^.

**Figure 1.**
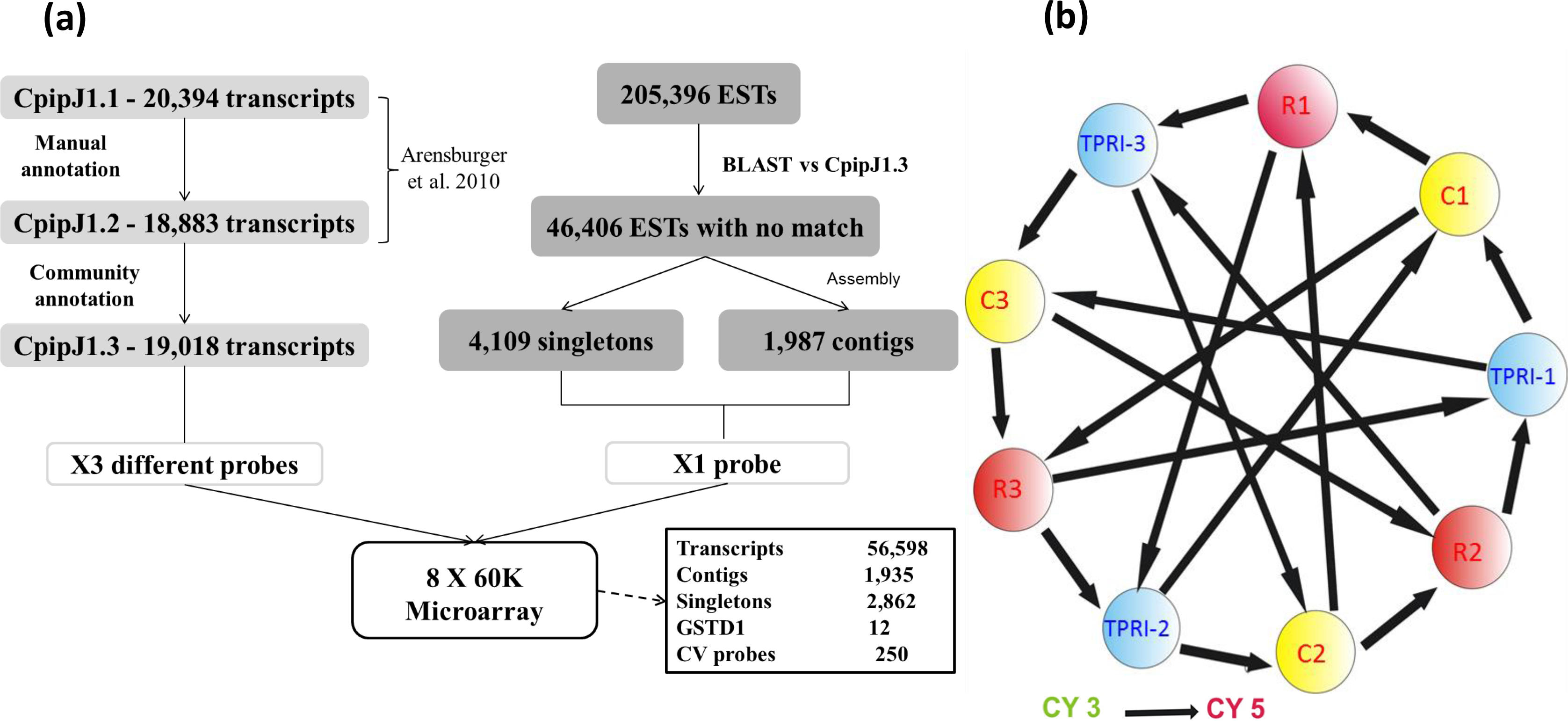
Overview of *Culex quinquefasciatus* whole-transcriptome analysis. A) Design of the 8 x 60K Agilent microarray. CpipJ1: consensus gene set of the automated gene prediction from the *C. quinquefasciatus* Johannesburg strain genome sequence. EST: expressed sequence tags. GSTD1: Glutathione S transferase D1. CV probe: coefficient of variation. B) Interwoven hybridization loop design for comparison between bendiocarb exposed and non-exposed Ugandan field-collected mosquitoes and the TPRI susceptible strain. Circles represent pools of 10 females. C: Uganda non-exposed mosquitoes (sympatric control), R: Uganda Resistant mosquitoes, TPRI (Tropical Pesticides Research Institute): *C. quinquefasciatus* susceptible strain from Tanzania.

**Figure 2.**
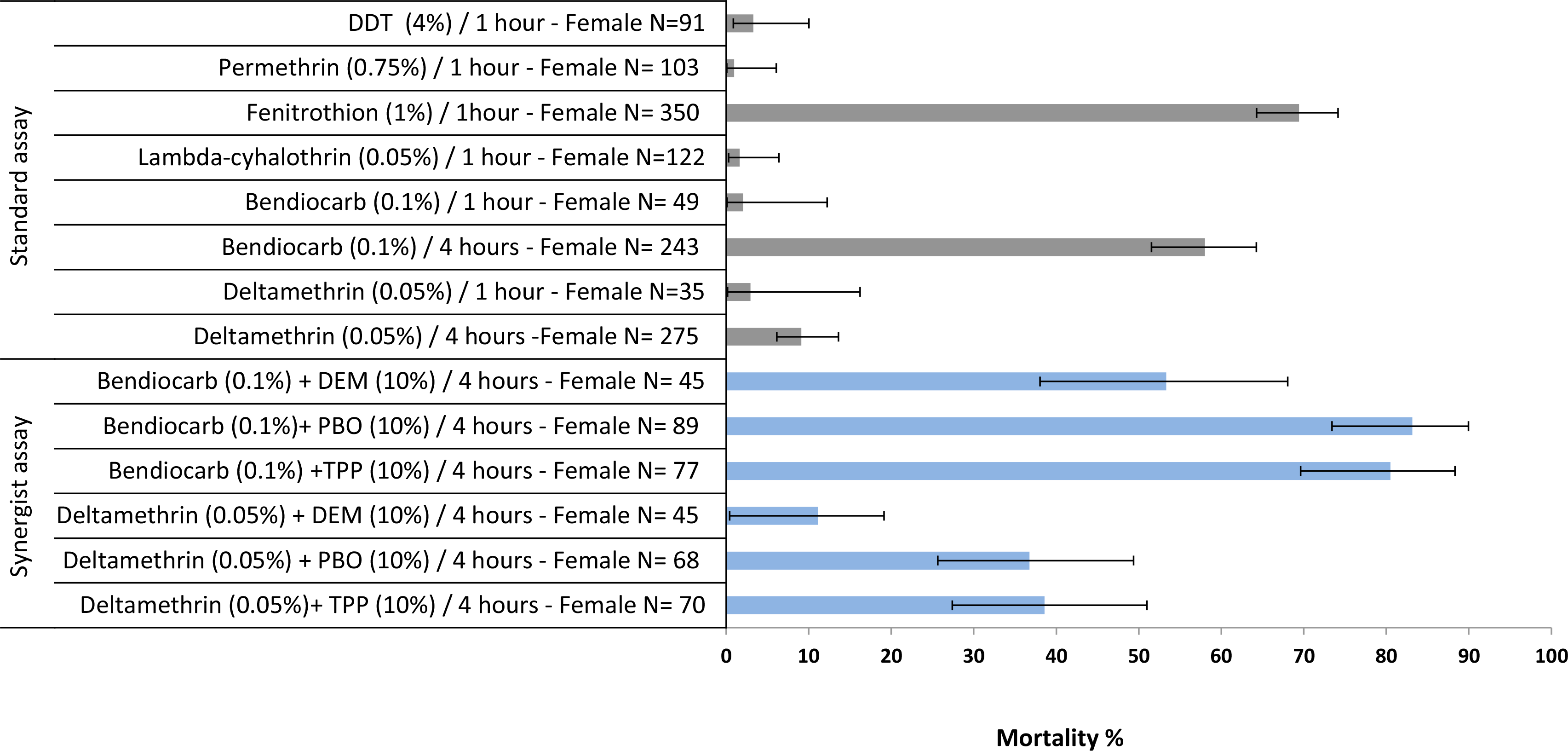
Insecticide susceptibility status of *C. quinquefasciatus* from Tororo (Uganda). Bioassay results following exposure to WHO insecticide treated papers at standard conditions and effect of insecticide synergists on the susceptibility status. Grey and blue bars represent WHO standard and synergists bioassay, respectively. Error bars represent 95% CI. PBO: piperonyl butoxide, DEM: diethyl maleate, TPP: triphenyl phosphate.

#### RNA extraction, labelling and Hybridization

Total RNA was isolated from three pools of 10 female mosquitoes for each group using the RNAqueous-4PCR kit (Ambion) according to the manufacturer’s instructions. Total RNA quantity was assessed using a Nanodrop spectrophotometer and RNA quality assessed on an Agilent Bioanalyzer. Each pool of RNA was individually labelled with Cyanine-3 and Cyanine-5 (Cy3 and Cy5) using the Low input Quick Amp Labelling Kit (Agilent Technologies) followed by purification through Qiagen RNeasy Columns (Qiagen) with quality and quantity checked using a Nanodrop and Bioanalyzer, respectively.

Before hybridization, 300 ng of Cy3 and Cy5 labeled cRNA was fragmented using the gene expression hybridization kit (Agilent) in a total volume of 25 μl including 5 μl of 10x blocking agent and 1 μl of 25x fragmentation buffer. The fragmentation reaction was incubated at 60°C for 30 min, then chilled on ice for 2 min before addition of 25 μl of 2 x GE hybridization buffer Hi-RPM. Each array was hybridized using 45 μl of the fragmentation solution for 17 hours at 65°C and 10 rpm. After hybridization, arrays were washed with wash buffers 1 and 2 for 1 min each, followed by acetonitrile for 10 sec and finally fixation solution for 30 sec. Arrays were scanned using the microarray scanner system (Agilent Technologies) and feature extraction performed using Feature Extraction software (Agilent Technologies) according to the manufacturer’s recommendations. All arrays passed the Agilent quality control with QC score ≥10.

### Data Analysis

All microarray data analysis was performed in R ^43^. Array normalization was carried out using the Limma 3.2.3 package ^44^ and data analysis performed using MAANOVA software ^45^ to detect overall differential levels of gene expression across the three treatment groups. The top overexpressed genes were selected after an ANOVA *F*-test based upon a false discovery rate (FDR) decision criteria of log_10_ (Q value) > 2.5. Within this subset of significantly differentially expressed probes, those that were significantly overexpressed in the Bendiocarb-exposed mosquitoes were pinpointed by comparing expression pattern in all pair-wise comparisons (see Table 1).

**Table 1.** Top differentially expressed genes from microarray analysis comparing Uganda resistant and sympatric controls compared to TPRI susceptible strain.

| Gene | Transcript | GO | log Fold-Change (-log Q value) |  |
| --- | --- | --- | --- | --- |
|  |  |  | Ugandan Bendiocarb vs TPRI | Ugandan Sympatric vs TPRI |
| CPIJ020018-RA | cytochrome P450 6Z16 | monooxygenase activity | 3.13 (2.80) | 2.59 (2.80) |
| CPIJ005900-RA | cytochrome P450 6N23 | monooxygenase activity | 2.13 (2.80) | 1.87 (2.80) |
| CPIJ003654-RA | electron transfer | electron carrier activity | 2.36 (2.58) | 2.23 (2.58) |
| CPIJ010823-RA | serine protease 27 precursor | serine-type endopeptidase activity | 4.41 (2.53) | 3.86 (2.44) |
| CPIJ013083-RA | brain chitinase and chia | hydrolase activity, hydrolyzing O-glycosyl compounds | 1.31 (2.44) | 0.93 (2.38) |
| CPIJ016928-RA | 15.4 kda salivary peptide | No information in VectorBase | 2.99 (2.44) | 2.73 (2.42) |
| CPIJ004984-RA | serine proteases 1/2 precursor | serine-type endopeptidase activity | 4.26 (2.44) | 3.60 (2.40) |
| CPIJ011746-RA | R2D2 | double-stranded RNA binding | 1.89 (2.41) | 1.72 (2.40) |
| CPIJ004019-RA | saccharopine dehydrogenase | oxidoreductase activity | 1.47 (2.44) | 1.05 (2.30) |
| CPIJ004019-RA | saccharopine dehydrogenase domain | oxidoreductase activity | 1.24 (2.40) | 1.11 (2.40) |
| CPIJ002104-RA | plasma alpha-L-fucosidase precursor | catalytic activity | 1.32 (2.44) | 0.90 (2.26) |
| CPIJ015009-RA | histone H3.3 type 2 | DNA binding | 1.32 (2.40) | 1.13 (2.38) |
| CPIJ000182-RA | N-acetylneuraminate lyase | catalytic activity | 2.44 (2.42) | 1.76 (2.26) |
| CPIJ011590-RA | conserved hypothetical protein | No information in VectorBase | 0.87 (2.37) | 0.81 (2.38) |
| CPIJ019290-RA | DNA ligase 4 | DNA binding | 1.22 (2.34) | 1.20 (2.35) |
| CPIJ016792-RA | hypothetical protein | No information in VectorBase | 3.39 (2.38) | 2.94 (2.29) |

Additionally, the pattern of differential transcript level was also characterized through pair-wise comparisons between Ugandan resistant, sympatric control and the TPRI strain using a Student’s t-test with genes considered differentially overexpressed where *P* < 0.05.

Functional characterization of differently expressed transcripts detected from the ANOVA and pair-wise comparison were then submitted to a Gene Ontology (GO) analysis to classify probes on their GO categories (cellular components, biological process and molecular functions). For this, a sub-set of overexpressed probes were selected based on the threshold of log_10_ (Q value) > 2 of the ANOVA analysis to capture a broader selection of highly overexpressed probes (see Fig. 4A). This was then submitted to VectorBase for functional annotation (GO term identification) using the Biomart tool. The REVIGO web server ^46^ was employed to summarize and visualize the distinct GO terms identified. Relatedness among GO terms was assessed using the uniqueness method, followed by clustering of GO terms with closer semantic similarity.

### RT-qPCR Validation

Reverse-transcription quantification PCR (RT-qPCR) was applied to confirm the expression profile of three out of 16 top candidate gene identified by the microarray (*CPIJ020018* [=*Cyp6z18*], *Cyp6n23, R2D2* Table 1) using two housekeeping genes - *40S ribosomal protein S3* and *β-tubulin* - as endogenous controls. Specific primers to amplify PCR fragments with size ranging from 107 to 176 bp (see Supplementary Table S1) were designed using primer3 software ^47^; however, only for four genes *Cyp6z18*, *R2D2*, *40S ribosomal protein S3* and *β-tubulin*) was it possible to design primers spanning exon junctions. Specificity of primer sets was verified by identification of a single symmetrical amplicon peak following melting curve analysis. Additionally, PCR efficiency was verified using a 10-fold serial dilution of standard cDNA with only primer sets with an efficiency ranging from 90 to 110% taken forward for RT-qPCR reactions.

Two technical replicates of all RT-qPCR reactions were carried out for each gene in a total volume of 20 μl including 1 μl of cDNA (1:4 stock diluted), 10 μl of Brilliant II SYBR^®^ master mix (Agilent Technologies) and 100 nm of each forward and reverse primer. Amplification was conducted under standard qPCR reaction conditions on the Mx3500P qPCR system (Agilent Technologies). Gene expression quantification of the three selected genes was assessed according to the ΔΔ^ct^ method^48^.

### Manual annotation of the *CPIJ020018* gene region

During qPCR primer design for the top candidate gene *CPIJ020018* (annotated as *Cyp6z16* in VectorBase CpipJ1.3 assembly #1, annotation 1.3), we concluded that the available, automated gene annotation of this particular gene was unreliable. The genomic sequence of this region in VectorBase includes a region of about 810 bp in supercontig 3.2948 with no nucleotide sequence information, which spanned the automated annotation of *Cyp6z16*. We suspected additional coding sequence to lie within this region. To confirm this, we designed primers to span the complete region and amplified a 4.7 kb region covering the full length of the candidate gene genomic sequence.

PCR reactions to amplify the *Cyp6z16* genomic region were conducted in a final volume of 20 μl including 40 ng of genomic DNA, 1 x Phusion HF buffer, 200 μM each dNTP, 0.5 μM each primer Cx_6Z16F and Cx_6Z16R (see Supplementary Table S2) and 0.02 U/μl Phusion DNA polymerase. Reaction conditions were 98°C for 30 sec, 30 cycles of 98°C for 10 sec, 62°C for 30 sec and 72°C for 3 min, with a final extension of 72°C for 5 min. PCR products were purified using the GeneJET PCR purification kit (Thermo Scientific) then cloned into pJET1.2 PCR vector (Thermo Scientific). Finally, nine primers (Supplementary Table S2) were used for sequencing the full length of cloned PCR product after plasmid purification using the GeneJET Plasmid Miniprep Kit (Thermo Scientific).

Sequence traces were analyzed using CodonCode Aligner version 4.2.2. Following removal of vector sequences a single contig was built from overlapping sequences. This contig sequence was then used for gene structure prediction and transcript annotation using the Augustus web interface ^49^.

### Transgenic expression in *Drosophila* flies

The full-length sequence of *Cyp6z18* was codon-optimised for *Drosophila* and synthesised with *Eco*RI-*Xba*I sites then cloned into the pUAST.attB vector (provided by Dr J Bischof, University of Zurich) by Genscript (Piscataway, NJ, USA). Transformation was performed using the PhiC31 system with plasmids injected into the germline of *D. melanogaster* with a chromosome 2 *attP* landing site (y w M(eGFP, vas-int, dmRFP)ZH-2A; P{CaryP}attP40) by the University of Cambridge Fly Facility. A single transgenic line was generated and balanced. To induce expression of *Cyp6z18*, flies were crossed to the Act5C-GAL4 strain (y1 w*; P (Act5C-GAL4-w) E1/CyO,1;2) (Bloomington Stock Center, IN, USA). For each treatment, *Cyp6z18* transgenic or untransformed controls, three crosses of 12-15 females to 6-7 Act5C-GAL4 males was performed. Shortly after pupae were seen, the parental generation was transferred into new rearing vials to continue egg laying. The date of appearance of newly-emerged offspring was observed for a total of 8 *Cyp6z18* and 11 control rearing vials.

## RESULTS

### Insecticide susceptibility status

Insecticide resistance levels following WHO susceptibility tests on F1 female mosquitoes to all six insecticides tested are shown in Figure 2. The lowest mortality (0.97%) was observed to permethrin while for DDT, lambdacyhalothrin, bendiocarb and deltamethrin the mortality rate ranged from 1.63-3.29% (Fig. 2). For fenitrothion we observed the highest mortality among the insecticides tested (69.42% 95% CI 64.27-74.16). To investigate the effect of exposure duration on mortality, bioassays with bendiocarb and deltamethrin were also carried out for four hours. For both insecticides, an increase in mortality was detected (Fig. 2); however, only for bendiocarb did the mortality increase significantly from 2.04 (95% CI 0.11-12.24) to 58.02% (51.53-64.25) whilst for deltamethrin the increase was non-significant (to 9.09%; 95% C.I. 6.12-13.59%).

Adult mosquitoes were also assayed with bendiocarb and deltamethrin for four hours exposure after pre-exposure in parallel to three synergist compounds (TPP, DEM and PBO). Synergism was observed for both insecticides following TPP and PBO pre-exposure whilst no significant effect on mortality was detected for DEM (Fig. 2). TPP and PBO significantly increased the mortality of bendiocarb from 58.02% (51.53-64.25) to 80.51% (69.6-88.34) and 83.14% (73.41-89.96) respectively.

### Frequency of *Ace-1* resistant alleles in bendiocarb selected mosquitoes

Both dead and alive mosquitoes following exposure to 0.1% bendiocarb (4h) were genotyped for the *Ace1*-119S mutation using a custom *Taq*man assay. The wild-type allele was observed at the highest frequency (86.22%; 95% CI 81.44-89.92%) with homozygous genotypes predominating (72.44%; 95% 66.64-77.57%). No homozygous resistant genotypes were detected (Fig. 3A). There was a highly significant association between the *Ace1*-119S allele and bendiocarb resistant phenotype (Fig. 3B; *P* < 0.0001) with an OR of 25 (3.37-186).

**Figure 3.**
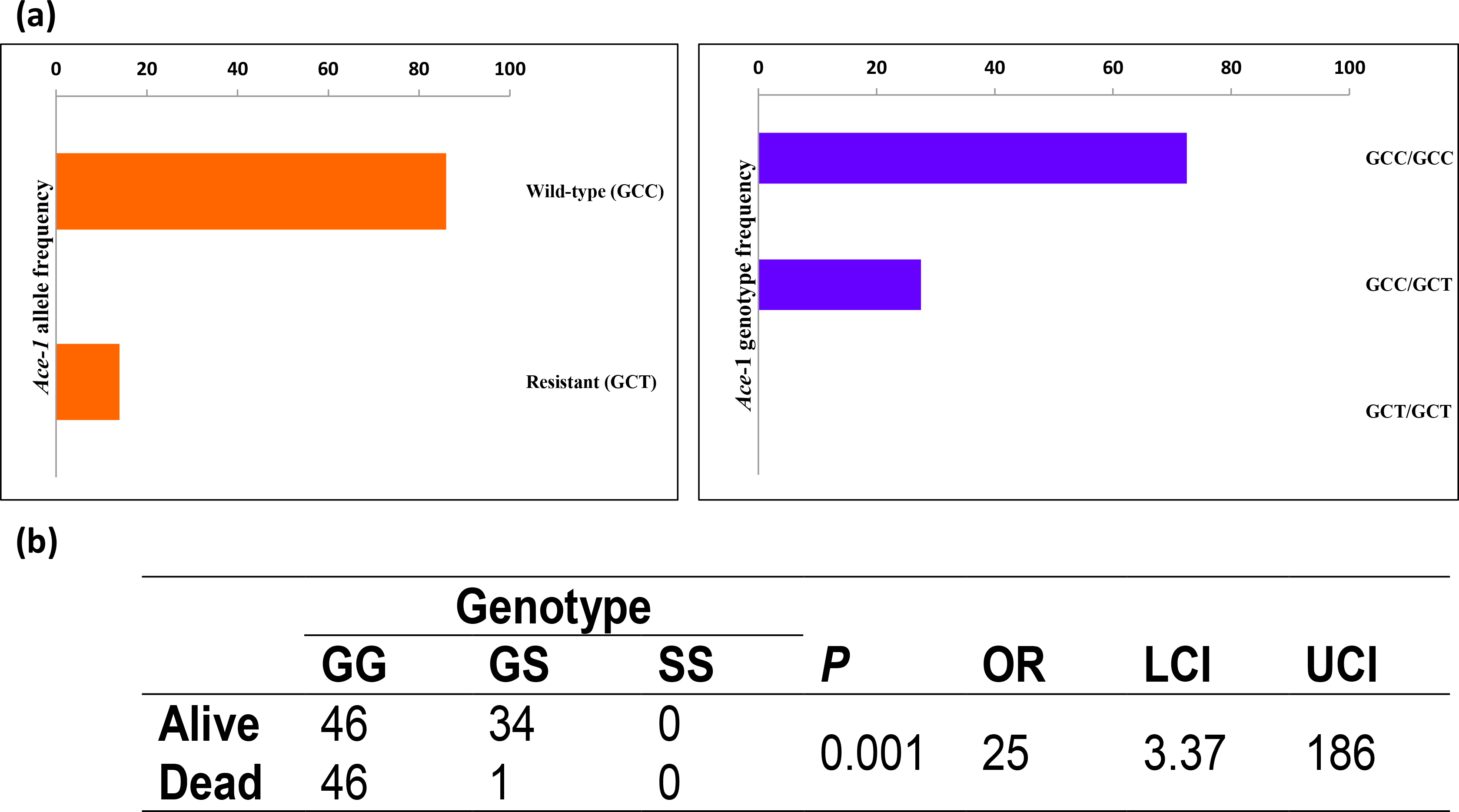
*Ace1-119S* allele and bendiocarb association test in *C. quinquefasciatus*. A) *Ace*-1 allelic and genotypic frequencies B) association of the *Ace*-1 genotype and bendiocarb (0.1%)/4 hours resistant phenotype.

### Gene expression profiling of bendiocarb selected mosquitoes

To identify candidate genes associated with resistance in bendiocarb selected mosquitoes, transcriptomic profiles of three groups of *Ace-1* wild-type (119G) mosquitoes were compared: Ugandan bendiocarb exposure survivors, Ugandan Control exposed (sympatric control) and a control exposed fully susceptible TPRI strain. Among the three groups we identified 32 probes significantly differently expressed in the ANOVA analysis applying a threshold of −log_10_ false-discovery rate (FDR) adjusted *P* value > 2.5, of which 50% of the probes had higher expression in the Ugandan exposed compared to both the sympatric control and TPRI (Table 1). The list of top candidate genes included two P450s: *CPIJ020018* (shown to be *Cyp6z18* following reannotation - see below) and *Cyp6n23*, which displayed increases in gene expression compared to the TPRI strain of 8.75 and 4.37 fold, respectively.

For functional characterization of the most significantly differentially expressed probes, a list of probes with a −log_10_ false-discovery rate (FDR) > 2 obtained from the ANOVA (Fig. 4A) comparison was submitted to Gene Ontology (GO) analysis. After removing duplicate probes, 358 unique genes were submitted to VectorBase for the GO term search using Biomart (for complete annotation see Supplementary Table S3.) In total 298 terms were obtained with the majority of extracted GO terms clustered on the molecular function and biological process categories (see Supplementary Table S4 for GO Terms’ frequencies and description). Among the top 10 enriched terms, GO linked to metabolic process, biosynthetic process, transport, membrane, integral component of membrane, nucleus, binding, cation binding and metal ion binding were observed with percentage of annotations ranging from 2.16 to 75.39% (Fig. 4B-D). Metabolic process was the predominant term across all categories corresponding to 75.39% of enriched GO terms, while in the category of molecular function and cellular component was observed a large proportion of terms associated with binding functions 21.23% and membrane 61.59%, respectively.

**Figure 4.**
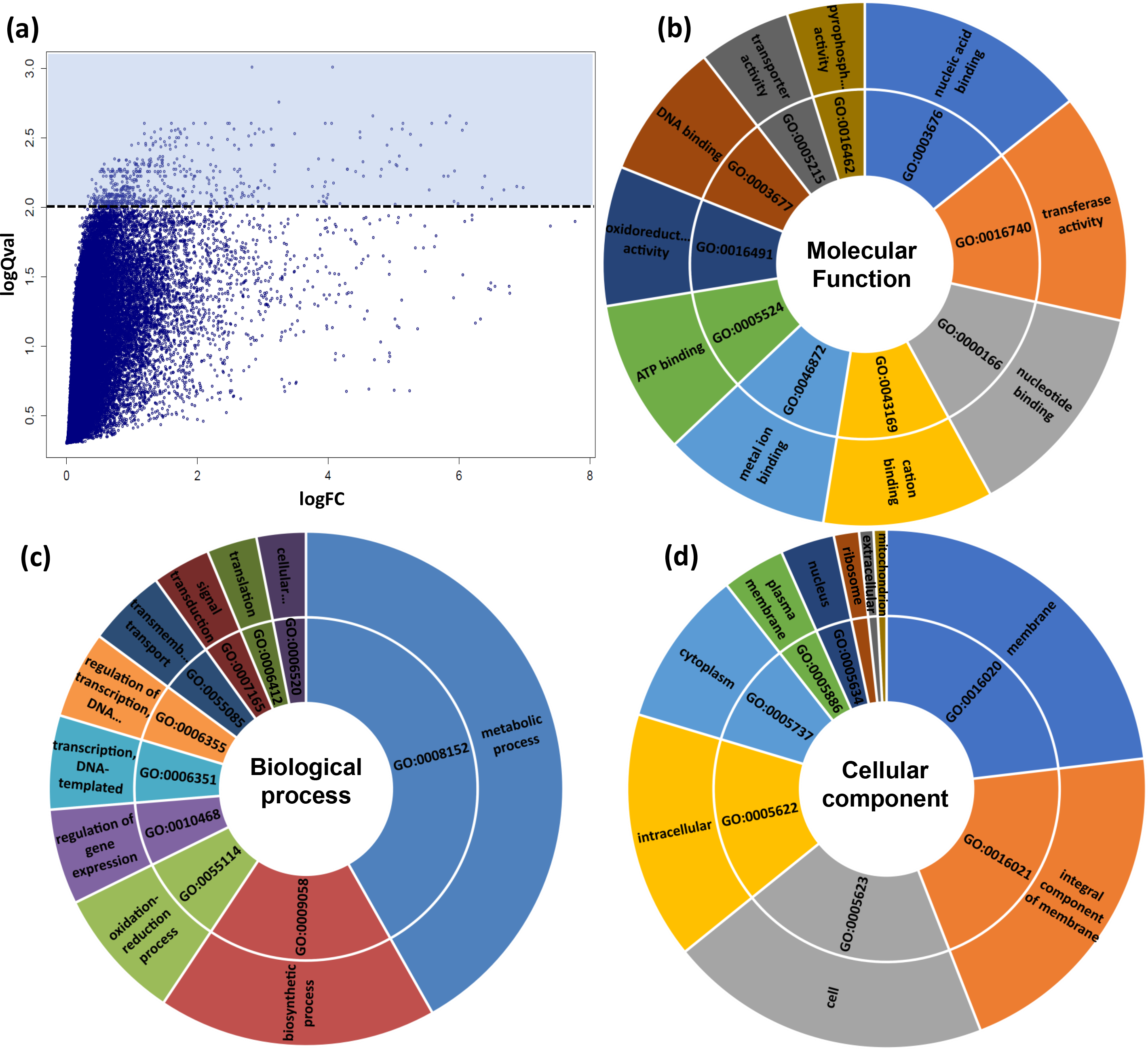
Candidate genes differentially transcribed in *C. quinquefasciatus* bendiocarb selected mosquitoes. A) Changes of gene expression between the three groups (Uganda exposed and un-exposed and TPRI) presented as a volcano plot. B, C and D are sunburst plots showing representative top 10 GO term clusters (molecular function, biological process and cellular component, respectively) of differentially expressed transcripts with FDR>2.0.

To explore further the transcriptomic profile of the bendiocarb resistant phenotype, pairwise comparison using Student’s t-test was applied to compare exposed and unexposed Ugandan mosquitoes to the TPRI susceptible strain. Significantly up and down-regulated genes with a fold-change > 2.5 were also investigated by GO analysis. For down-regulated genes we observed similar figures between exposed and unexposed mosquitoes in contrast to up-regulated genes where we identified 33 genes exclusively in the exposed mosquitoes (Fig. 5A). Pairwise comparisons also identified eight significantly expressed genes putatively associated with insecticide detoxification: three exclusively in the pools of exposed mosquitoes: GSTs (CPIJ018629-RA; fold-change 2.79 and CIPJ018632-RA, fold-change 2.0) and esterase (CPIJ013918-RA; fold-change 2.86) whereas for the other five: P450 (CPIJ020018-RA), GSTs (CPIJ010814-RA, CPIJ018624-RA, CPIJ018626-RA) and esterase (CPIJ013917-RA) were observed in both Ugandan exposed and unexposed mosquitoes.

**Figure 5.**
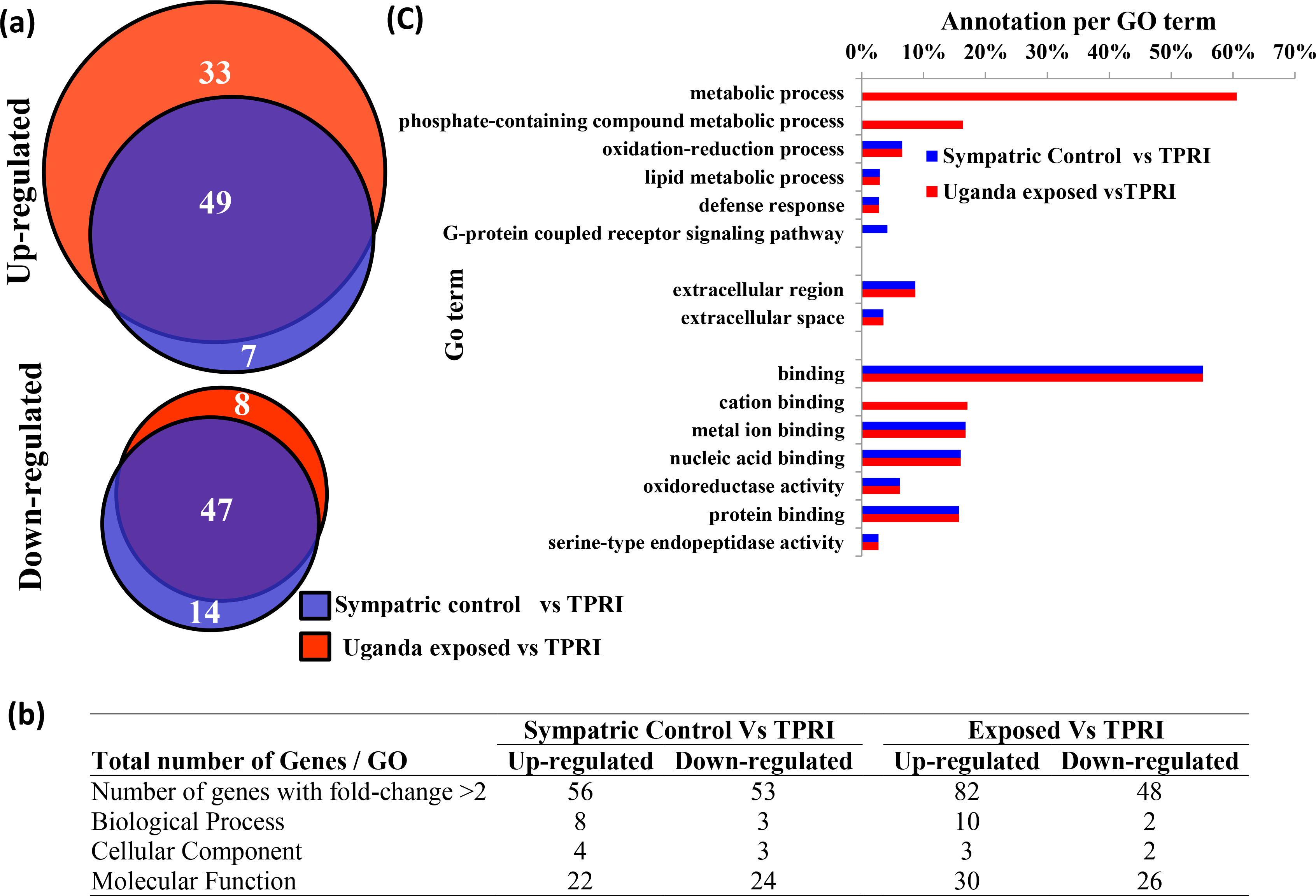
Transcriptomic profile of differentially expressed genes with fold-change >2 in Uganda exposed and sympatric mosquitoes compared to TPRI. A) Venn diagram showing the overlap of up- and down-regulated transcripts between the three groups. B) Comparison of the number of GO terms identified by each pair-wise comparison. C) GO term enrichment of up-regulated transcripts between the groups with frequency higher than 2%.

In general, the GO term enrichment of the up and down-regulated genes for the Ugandan exposed and unexposed mosquitoes shows a similar list of terms (Fig. 5B) with the exception of three terms: metabolic process, phosphate-containing compound metabolic process and cation biding that were observed exclusively on the exposed mosquitoes with frequencies of 60.61, 16.39 and 17.07%, respectively (Fig. 5C). Nevertheless, this analysis excluded 39 and 18 up and down-transcribed genes (See Supplementary Table S5 and S6), respectively, such as esterases (CPIJ013917-RA and CPIJ013918-RA) and Heat shock proteins (CPIJ013880-RA and CPIJ005642) for instance, that were not associated with GO terms from VectorBase.

### Candidate gene validation by qPCR

The expression patterns of three candidate genes, randomly chosen from the list of top candidates, (both P450 genes *Cyp6z18* and *Cyp6n23*, plus *R2D2*) were additionally assessed by qPCR. Satisfactory PCR efficiency ranging from 91.4 to 101.8%, (within the 10% acceptable variation) was obtained for primer pairs designed for candidate genes and endogenous controls (Supplementary Fig. S1). Additionally, the primer sets were specific, with a single symmetrical amplicon peak in melting curve analyses (Supplementary Fig. S2). For all three candidate genes (*Cy6z17*, *Cyp6n23* and *R2D2*) we observed a good correlation between the microarray and RT-qPCR expression fold-change with ratios of up-transcription level detected by both methods differing by less than 1.5x (Supplementary Fig. S3).

### Annotation of *CPIJ020018*

After identification of *CPIJ020018* as the top candidate gene in the microarray results, further analyses on the genomic and cDNA sequences available from VectorBase were conducted. This revealed an atypical gene architecture for a P450 gene with the presence of one intron > 3Kb (Fig. 6A). Closer analysis of the genomic sequence indicated a region of 809 bp with no nucleotide information (Ns) internal to the *CPIJ020018* genomic sequence. *In silico* analysis of a contig constructed after PCR amplification and cloning of the complete region encompassing the genomic region of interest (accession number MH822866), suggested the presence of two distinct P450s instead of the single *Cyp6z16* predicted by the automated annotation in VectorBase. The two genes predicted are each composed of two exons separated by one intron (Fig. 6B and Supplementary material 1). BLAST analysis of both predicted amino-acid sequences against *Anopheles gambiae* and *Aedes aegypti* sequences available on VectorBase and the Cytochrome P450 homepage ^50^ shows the top hits belong to CYP genes from the Z family. Following submission to the P450 nomenclature database the partial P450 is now labelled *cyp6z16* and the full-length novel P450, which is interrogated by the microarray probes (hitting *CPIJ020018* exon-2) is formally labelled *Cyp6z18* (note *Cyp6z17* is annotated on supercontig 3.3058). Together the gene prediction and BLAST analysis carried out indicate that the top candidate gene from the microarray analysis is *cyp6z18* and not *cyp6z16* (which is not interrogated by any probes due to incorrect annotation). *cyp6z18* is closely related to other P450s which have previously been associated with metabolic resistance in both *An. gambiae* and *Ae.aegypti* mosquitoes (Fig. 6C). We note that whilst our microarray design was undertaken on the CpipJ1 assembly, the CpipJ2 assembly (available from April 2014) did not revise the assembly or annotation of this scaffold.

**Figure 6.**
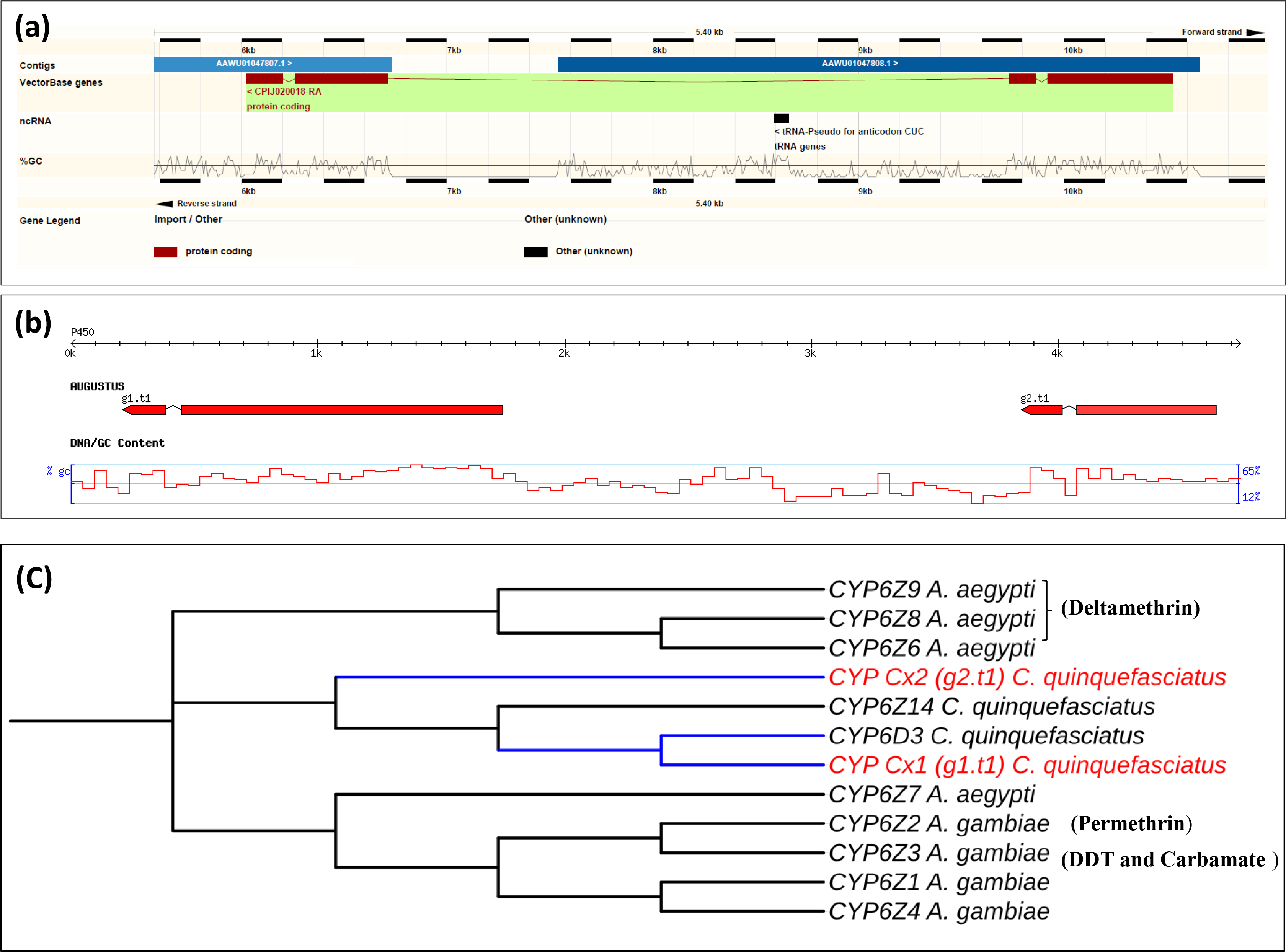
*Cyp6z16* and *Cyp6z18* predicted gene structure and annotation. A) output of the VectorBase genome Browser suggesting a gene architecture with four exons and three introns. B) Schematic representation of *CPIJ020018* after re-annotation using Augustus software, indicating two distinct genes here named *Cyp6z18* (g1.t1) and *Cyp6z16* (g2.t1). C) Unrooted distance neighbour joining tree showing phylogenetic relationship of the predicted gene *Cyp6z18* from *C. quinquefasciatus* to *Aedes aegypt* and *An. gambiae* cytochrome P450s from the CYP6 gene family. The percentage of bootstrap confidence values from 1000 replicates is shown at the nodes.

### Transgenic expression of *cyp6z18* in *Drosophila* flies

*Drosophila melanogaster* flies were transformed with a transgene encoding *cyp6z18* for expression using the GAL4/UAS system. Individuals containing the *cyp6z18* gene and untransformed controls were crossed to flies of the Act5C-GAL4 driver line to induce ubiquitous expression of *cyp6z18*. Expression of *cyp6z18* negatively impacted fly fitness, evident in prolonged pre-adult development. The number of days between placing adult flies in a rearing vial and the first offspring emerging was measured. A significant difference in this value was detected, with an average of 23 days for *cyp6z18*-expressing flies (95% CI 21.21-25.04) and 16 days for untransformed control flies (95% CI 15.23-16.41) (unpaired t-test with Welch’s correction *P* < 0.0001, *N* = 19). Because of this fitness cost, experiments to test the impact of *cyp6z18* expression upon insecticide resistance were not pursued.

## DISCUSSION

Insecticide resistance is widespread in malaria vectors in Uganda ^35,51–55^ and known to result from both target-site and detoxification mechanisms ^55–57^. In this study, we investigated the insecticide susceptibility status and possible mechanisms involved with insecticide resistance in the LF vector *C. quinquefasciatus* mosquitoes collected in a region of Uganda (Tororo) where both *Anopheles gambiae* and *Anopheles funestus* exhibit resistance to the insecticides currently used for malaria control ^35,54,55^. We have shown previously ^39^ that the pattern of target-site mutations in Ugandan *Culex* varies regionally despite intense gene flow, suggesting a heterogeneous pattern of insecticidal selection pressures. Here we show that in *Culex* from Tororo, eastern Uganda, resistance to three of the four classes of insecticide occurs at high levels and, at least for bendiocarb, is mediated by both target-site and metabolic mechanisms. Although, the G119S mutation in *Ace-1* is not at high frequency, it is strongly associated with resistance with an OR of 25 (95% CI 3.4-186). With such a strong resistance association it seems surprising that the 119S allele is at such low frequency (14%9 5% CI 7.9-22.4%) in the Nagongera population. This is a much lower frequency than observed in other vector mosquitoes where carbamates are routinely used in vector control ^58^. Whilst insecticides linked to selection of *A*ce-*1* resistant alleles have not been officially applied for vector control in the studied area for over one decade ^59,60^ indoor residual spraying (IRS) using bendiocarb was conducted in Tororo from December 2014 - February 2015 ^61,62^, after the period of collection of these samples, and hence this allele frequency is higher than expected. Nevertheless, an indirect source of insecticide exposure due to agricultural activities could also be shaping the insecticide resistance as described for *C. quinquefasciatus* populations from Iran ^63^. However, the *Ace-1* mutation is known to confer a fitness disadvantage which may explain the low frequency of the resistance associated allele. This recent evolution of insecticide resistance in Ugandan *Culex* populations, possibly in response to malaria vector control, has also been suggested in other African countries such as Tanzania and Zambia ^15,64^, where moderate resistance has also been detected to bendiocarb, in contrast to the high levels of resistance detected to both pyrethroids and DDT. The potential involvement of metabolic resistance in bendiocarb resistant *C. quinquefasciatus* from Tororo is supported by the synergist assays, which show an increase in mortality to bendiocarb when mosquitoes were pre-exposed to either TPP (a synergist of esterases) or PBO (a synergist of cytochrome P450s). Together the *Ace-1* genotyping and synergist assay data strongly suggested an alternative mechanism of resistance to the well-known *Ace1*-119S target-site mutation in the Ugandan bendiocarb resistant phenotype. Transcriptomic profiling of *Ace-1* wild-type mosquitoes identified two P450s (*CPIJ020018* and *Cyp6n23*) with the highest up-regulation in the resistant samples among the top candidate genes. Over-expression of both these cytochrome P450s (*CPIJ020018* and *cyp6n23*) identified in our analysis are especially relevant as many genes belonging to this gene family have been associated with insecticide metabolism in a variety of vector species ^26,65,66^. The likely association of this candidate cytochrome P450s with the carbamate resistant phenotype is supported also by the synergism effect of the P450 inhibitor PBO ^67^. Most CYPs previously associated with the insecticide resistance phenotype in mosquitoes, (e.g. *cyp6p3*, *cyp9j32* and *cyp6m10*) are typically associated with metabolism of pyrethroids and DDT ^66,68,69^, while so far very few examples of bendiocarb metabolism by P450 have been reported. Recently, Edi *et al*. ^70^ demonstrated that *cyp6p3* is associated with the bendiocarb resistant phenotype in *An. gambiae* from Tiassalé Cote d’Ivoire and confirmed its capability to metabolise bendiocarb. Additionally, two other *An. gambiae* P450s (*cyp6z1* and *cyp6z2*) have also been demonstrated to be capable of metabolizing the carbamate insecticide carbaryl ^71^ and upregulation of the *An. funestus cyp6z1* is associated with carbamate resistance in this species ^72^.

Interestingly, both cytochrome P450s identified here belong to the CYP6 family, which includes most of the CYPs genes already described as insecticide metabolizers ^21^.

Further analysis of the genomic and transcriptomic sequences of the top candidate gene *CPIJ020018* (annotated as *cyp6z16* in VectorBase) suggests annotation inaccuracy. After re-annotation, our analysis indicates that *CPIJ020018* consists of two distinct CYP genes (*cyp6z16* and *cyp6z18*) instead of one as suggested by the automated annotation of VectorBase. Annotation problems as observed for *cyp6z16* may potentially occur for other genes (see ^73^ for annotation deficiencies of recent *Anopheles* sequencing), thus a review of *C. quinquefasciatus* gene annotation, especially of gene families linked to insecticide resistance, would benefit future transcriptomic monitoring of insecticide resistance. Whilst, the increased usage of RNASeq for differential gene expression analyses may suggest this is unnecessary, the typical RNA-Seq pipeline maps reads to annotated gene sets and hence a robust annotation of these gene families would still be beneficial. *C. quinquefasciatus* remains the poor relation of disease vectors with respect to assembly and annotation of genome sequence. Whilst the genome sequences of *A. gambiae* and *A. aegypti* have received important and successful genome updates ^74,75^ *Culex* remains assembled with a high number of scaffolds and hence gaps.

This microarray study also identified that four other detoxification genes (three glutathione S-transferases (GSTs) and one esterase) were also over-transcribed compared to TPRI, although they were not differently expressed between Tororo resistant and sympatric unexposed controls. High expression of these genes alone or in combination with other detoxification enzymes could also be a possible mechanism associated with the bendiocarb phenotype as reported previously^76,77^, although in our data no significant synergist effect by DEM, a GST inhibitor, was observed. Further characterization of these genes is warranted.

Although *C. quinquefasciatus* is a vector of important neglected tropical diseases, planning and management of vector control strategies for *Anopheles* species has received considerably more attention and funding. Whilst this could reflect that MDA is currently the primary intervention for LF eradication with vector control deemed secondary, a reduction of LF transmission by *Culex* species due to control programs aimed at anopheline species has been demonstrated ^78,79^.

Thus, in a possible integrated vector-control scenario the monitoring of insecticide resistance and determination of the mechanisms resistance for vector species of apparent secondary interest can be critical to effective integration ^80^.

## CONCLUSIONS

Our data demonstrate that although Ugandan *C. quinquefasciatus* mosquitoes had not been covered by a specific, targeted local vector control program, high levels of insecticide resistance were identified in the studied population, indicating that application of insecticide to control other species with public health importance such as anophelines through indoor residual spray (IRS) and insecticide treated nets (ITNs), or through application of insecticides for agricultural purposes could be driving the evolution of insecticide resistance in this population. This study also provides strong evidence that a metabolic mechanism is associated with the bendiocarb resistant phenotype observed in Tororo. Lastly, by a whole-transcriptomic analysis we identify two new candidate genes belonging to the cytochrome CPY6 gene family associated with metabolic resistance to bendiocarb.

## ACKNOWLEDGEMENTS

Sample collection was supported by Award Number U19AI089674 from the National Institute of Allergy and Infectious Diseases (NIAID). Genomic and transcriptomic studies were supported by Award Number BEX-6190/10-3 from Coordination for the Improvement of Higher Education Personnel (CAPES). WFSM is a Wellcome Trust Fellow - Training Fellowship in Public Health and Tropical Medicine (Reference number - 209305/Z/17/Z). KM was funded by the National Institutes of Health/National Institute for Allergy and Infectious Diseases (grant numbers HHSN266200400039C; HHSN272200900039C).

We thank Dr Johannes Bischof, Institute of Molecular Life Sciences, University of Zurich for the plasmid.

## CONTRIBUTIONS

W.F.S.M., C.S.W. and M.J.D. conceived and designed the experiments. W.F.S.M., E. J. R., C.S.W., A. T. I. and D. P. performed the experiments. C.S.W and K. M. designed the array. W.F.S.M., C.S.W. and M.J.D. analysed the data. C.S.W, M.J.D. and H. D. M. Contributed reagents/materials/sample collections. W.F.S.M. wrote the paper. C.S.W and M.J.D., reviewed the final manuscript.

The authors have declared no conflict of interest

