## Supplementary Figure for "Transcriptomic analysis of insecticide resistance in the lymphatic filariasis vector *Culex quinquefasciatus*"

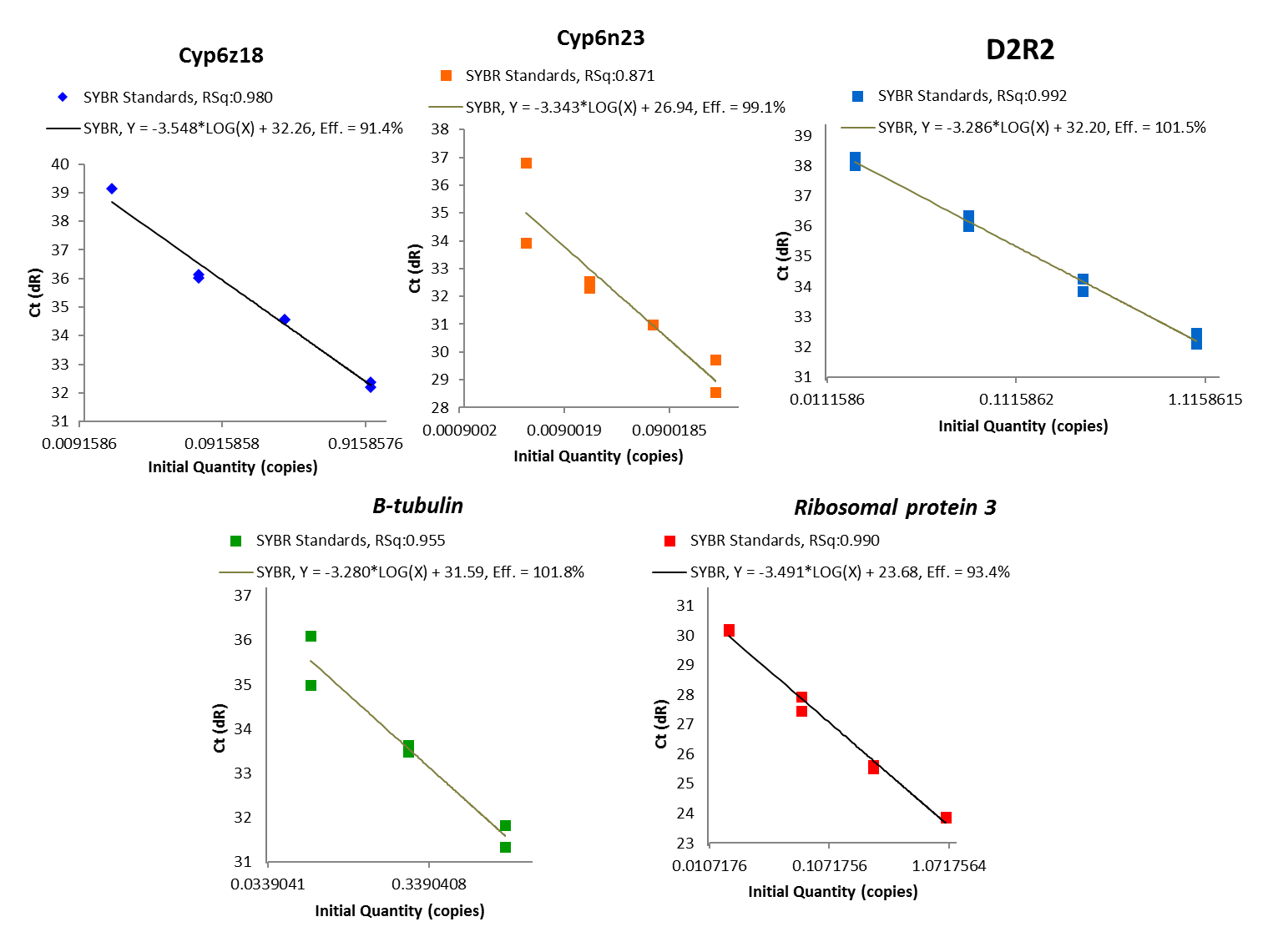

**Figure S1**: Standard curve from primers used on real-time PCR for microarray candidate genes validation. Squares correspond to 1x10 serial dilution of cDNA. qPCR efficiency (Eff) and coefficient of determination (Rsq) were calculated for each prime primer based two replicated.

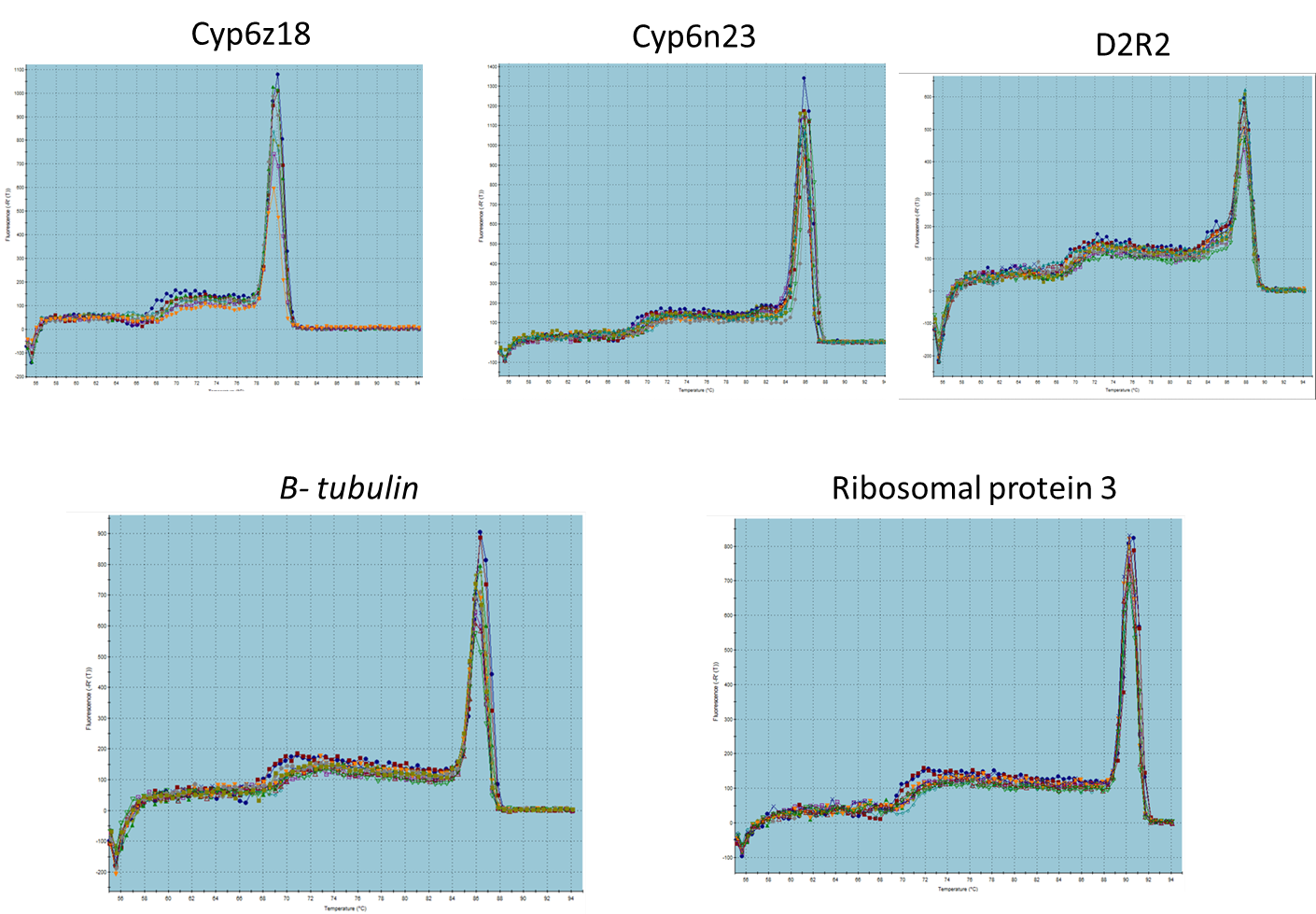

**Figure S2:** Validation of qPCR primers. Dissociation curves of real-time PCR amplification of microarray top candidate genes and endogenous control.

Log 2-fold change

| Selected genes from microarray data | qPCR | microarray |
| --- | --- | --- |
| Cyp6n23 Exposed vs TPRI | 3.63085 | 2.13 |
| Cyp6n23 Control Vs TPRI | 2.57687 | 1.87 |
| D2R2 Exposed vs TPRI | 1.5969 | 1.89 |
| D2R2 Control Vs TPRI | 0.611291 | 1.72 |
| Cyp6z18 Exposed vs TPRI | 3.703 | 3.13 |
| Cyp6z18 Control Vs TPRI | 3.539 | 2.59 |

Figure S3: Pairwise comparison of microarray and qPCR data based on gene expression profile for three genes found to be significantly different expressed between Uganda exposed and non-exposed mosquitoes (sympatric control) and TPRI susceptible strain
