## Supplementary Material 1 for "Transcriptomic analysis of insecticide resistance in the lymphatic filariasis vector *Culex quinquefasciatus*"

```
# This output was generated with AUGUSTUS (version 2.7).
# AUGUSTUS is a gene prediction tool for eukaryotes written by Mario Stanke

# and Oliver Keller.
# Please cite: Mario Stanke, Mark Diekhans, Robert Baertsch, David Haussler
(2008),
# Using native and syntenically mapped cDNA alignments to improve de novo gene
finding
# Bioinformatics 24: 637-644, doi 10.1093/bioinformatics/btn013
# No extrinsic information on sequences given.
# Initialising the parameters ...
# aedes version. Using default transition matrix.
# Looks like /data/www/augustus/tmp/AUG-166261241/input.fa is in fasta format.
# We have hints for 0 sequences and for 0 of the sequences in the input set.
#
# ----- prediction on sequence number 1 (length = 4735, name = P450) -----
#
# Constraints/Hints:
# (none)
# Predicted genes for sequence number 1 on both strands
# start gene g1
P450    AUGUSTUS        gene    209     1747    0.94    -       .       g1
P450    AUGUSTUS        transcript      209     1747    0.94    -       .
g1.t1
P450    AUGUSTUS        stop_codon      209     211     .       -       0
transcript_id "g1.t1"; gene_id "g1";
P450    AUGUSTUS        terminal        209     381     0.98    -       2
transcript_id "g1.t1"; gene_id "g1";
P450    AUGUSTUS        initial 445     1747    0.96    -       0
transcript_id "g1.t1"; gene_id "g1";
P450    AUGUSTUS        intron  382     444     1       -       .
transcript_id "g1.t1"; gene_id "g1";
P450    AUGUSTUS        CDS     209     381     0.98    -       2
transcript_id "g1.t1"; gene_id "g1";
P450    AUGUSTUS        CDS     445     1747    0.96    -       0
transcript_id "g1.t1"; gene_id "g1";
P450    AUGUSTUS        start_codon     1745    1747    .       -       0
transcript_id "g1.t1"; gene_id "g1";
# coding sequence =
[atgatcatttactcgctgcttctcatcggtacctcgatctacctgatcctccggtacatctactcgtactgggatcgcc
#
atggcctgccgaacctcaaaccggacatcccgttcgggaacatccgtgccgtcgccctcaagcaggaatcgttcggcgtc
gccctgaacgctctccac
#
gccaaaaccacaggccaactggtcggaatctacctactcttccgcccggcgatcctaatccgggacgcccacctagccca
ccgcatcataacgtccga
#
cttcaactacttccacgaccgcggtgtgcactgtgacgagagctcggatcccttttcggcgcacctgtttgccctgcccg
ggaagaggtggcgcagcc
#
tgaggaacaagctgactccgaccttcaccgctgggcagctgcgcggtatgctgccgacgatcttggccgttgggaggaag
tttcagggtttcttggaa
```

#
cccaaggcgaagcgaggggaggtgattgaggccagggatttgatatcgcgctttgtgctggagatcgtggcgtcggtgtt
tttcggttacgagatcaa
#
ctcgattcacgatccgcaggattcgtttcggacggttttacgatcatttcgggaggacaatcacgtcacaaatttgagaa
cggttggtgcgtttttgt
#
gtccgacgttgctgaaggtcagccgggtcaaaacggtacctgaggtggtggacaattttgtaaacaaaatcattagggag
cagatcgagtttcgcgag
#
aagaacaacgttacgaggaaggatttcatccaactgttgatcgatctccgacgggagaagagtgatttcggactttcgtt
ggaacagtgcgcggccaa
#
tgtgtttttgttctacgtagcaggagcggatacctcaacggatgccatcacctacacggtccacgagctgacccatcgac
cggatcttatgaagaagg
#
ttcaagcagagatcgacgatgcgcttgagaagtccaacggtgaaatcaactacgatgtactccacgaaatgaaactttta
gacaactgcgtgaaggaa
#
acccttagaaagtacccgttcccaattttaaatcgcgagtgtacccaggactatcaggttccagattcgaagctgatcat
caggaagggaactccggt
#
gatcataccgctgcaagcgttcggaatgagtgaggagtacttcccggaacctaatcgctacctacctgaacgatttgatt
catccacaaagaattacg
#
acgaaaaagcttacattccatttggagatggtccgaggaattgtattggttcccggatgggcagtgccgtttcgaagatc
ggcatcatcatgctgctt
#
tcgaagttcaactttgaggcgactcaaggtgcggagataggctttgctcgggcccaaattgcgctggctccggagaatgg
catttcacttaagatttc
### caacaggataagaaactcgatataa]
### protein sequence =
[MIIYSLLLIGTSIYLILRYIYSYWDRHGLPNLKPDIPFGNIRAVALKQESFGVALNALHAKTTGQLVGIYLLFRPAIL
#
IRDAHLAHRIITSDFNYFHDRGVHCDESSDPFSAHLFALPGKRWRSLRNKLTPTFTAGQLRGMLPTILAVGRKFQGFLEP
KAKRGEVIEARDLISRFV
#
LEIVASVFFGYEINSIHDPQDSFRTVLRSFREDNHVTNLRTVGAFLCPTLLKVSRVKTVPEVVDNFVNKIIREQIEFREK
NNVTRKDFIQLLIDLRRE
#
KSDFGLSLEQCAANVFLFYVAGADTSTDAITYTVHELTHRPDLMKKVQAEIDDALEKSNGEINYDVLHEMKLLDNCVKET
LRKYPFPILNRECTQDYQ
#
VPDSKLIIRKGTPVIIPLQAFGMSEEYFPEPNRYLPERFDSSTKNYDEKAYIPFGDGPRNCIGSRMGSAVSKIGIIMLLS
KFNFEATQGAEIGFARAQ
### IALAPENGISLKISNRIRNSI]
### end gene g1
###
### start gene g2
P450 AUGUSTUS gene 3848 4639 0.69 - . g2
P450 AUGUSTUS transcript 3848 4639 0.69 - .
g2.t1

```
P450	AUGUSTUS	stop_codon	3848	3850	.	-	0
transcript_id "g2.t1"; gene_id "g2";
P450	AUGUSTUS	terminal	3848	4014	0.96	-	2
transcript_id "g2.t1"; gene_id "g2";
P450	AUGUSTUS	initial	4072	4639	0.73	-	0
transcript_id "g2.t1"; gene_id "g2";
P450	AUGUSTUS	intron	4015	4071	0.99	-	.
transcript_id "g2.t1"; gene_id "g2";
P450	AUGUSTUS	CDS	3848	4014	0.96	-	2
transcript_id "g2.t1"; gene_id "g2";
P450	AUGUSTUS	CDS	4072	4639	0.73	-	0
transcript_id "g2.t1"; gene_id "g2";
P450	AUGUSTUS	start_codon	4637	4639	.	-	0
transcript_id "g2.t1"; gene_id "g2";
# coding sequence =
[atgaccaacctagcgacgcagcagatgaagtatcgtgagaagaatgacttggctagaaaggatttcttgcagttgctga
#
atgatcttcaccaagttgatttgtcagctgaagagtgcgcatcgaatgtgaatctgttctacactgcaggttcggaaacc
accaaatctacagtcatc
#
tatactcttcacgaactagctcaccatccagaagttatgagacggcttgtggaagaagttgatgaatacgtcaagcaatc
aggtggtgagattagcta
#
cgatcttgttaagagtatgccatatttggacctgtgcgtgaaggaaaccctgagaaagtatcccggactgtttttcctga
accggaagtgcacccacg
#
actataaggttcccaactctcggctggtcatcaaaaagggtacccaaataattatcccgtcgatggcctacggcatggat
gagcggtgtttcccgaat
#
ccggagagctacatccccgaacgatttcttgaggagaccaaaaattacgacgaggacgcctacgcaccgtttggagaagg
accgcggaagtgtatcgc
#
tcctcgaatgggaattttcgtcgccaaagtgaccttggtgaggctgctgtccaagtttcggttcgaggctacgcaagagc
tgaaggttgagtttgccc
# cctcggtgattccgctcgtgccgaaggatggagtcaggatgaagattcacaaaagaagtgtttggtaa]
# protein sequence =
[MTNLATQQMKYREKNDLARKDFLQLLNDLHQVDLSAEECASNVNLFYTAGSETTKSTVIYTLHELAHHPEVMRRLVEE
#
VDEYVKQSGGEISYDLVKSMPYLDLCVKETLRKYPGLFFLNRKCTHDYKVPNSRLVIKKGTQIIIPSMAYGMDERCFPNP
ESYIPERFLEETKNYDED
# AYAPFGEGPRKCIAPRMGIFVAKVTLVRLLSKFRFEATQELKVEFAPSVIPLVPKDGVRMKIHKRSVW]
# end gene g2
###
```
